# RACER: A data visualization strategy for exploring multiple genetic associations

**DOI:** 10.1101/495366

**Authors:** Olivia L. Sabik, Charles R. Farber

**Affiliations:** Center for Public Health Genomics and Department of Biochemistry and Molecular Genetics, School of Medicine, University of Virginia, Charlottesville, VA 22908; Center for Public Health Genomics and Departments of Public Health Science and Biochemistry and Molecular Genetics, School of Medicine, University of Virginia, Charlottesville, VA 22908

## Abstract

Genome-wide association studies (GWASs) have identified thousands of loci associated with risk of various diseases; however, the genes responsible for the majority of loci have not been identified. One means of uncovering potential causal genes is the identification of expression quantitative trait loci (eQTL) that colocalize with disease loci. Statistical methods have been developed to assess the likelihood that two associations (e.g. disease locus and eQTL) share a common causal variant, however, visualization of the two loci is often a crucial step in determining if a locus is pleiotropic. While the current convention is to plot two associations side-by-side, it is difficult to compare across two x-axes, even if they are identical. Thus, we have developed the Regional Association ComparER (RACER) package, which creates “mirror plots”, in which the two associations are plotted on a shared x-axis. Mirror plots provide an effective tool for the visual exploration and presentation of the relationship between two genetic associations.

**Availability and Implementation:** RACER is provided under the GNU General Public License version 3 (GPL-3.0). Source code is available at https://github.com/oliviasabik/RACER.

**Supplementary information:** Supplementary data are available online with the paper, see the Supplemental Data Manifest.

## I. Introduction

Genome-wide association studies (GWASs) have identified thousands of loci associated with disease risk; however, the genes responsible for the majority of these disease-associated loci remain largely unknown (Gallagher and Chen-Plotkin, 2018). A common approach to identify causal genes is to determine if disease-associated variants also influence molecular phenotypes, such as gene expression (Nicolae *et al*., 2010). This approach has become more widely implemented as expression quantitative trait loci (eQTL) across many tissues have become available from projects such as the Genotype-Tissue Expression Project (GTEx) (GTEx Consortium, 2013). Several statistical approaches that provide formal evidence of colocalization between two associations (e.g. a disease locus and eQTL) have been developed (Giambartolomei *et al*., 2018; Hormozdiari *et al*., 2016; Nica *et al*., 2010, https://eithub.com/Ksieber/piccolo): however, effective visualization is often an important component of colocalization analyses to ensure the presence of a single pleiotropic association. A common convention is to plot two associations separately using LocusZoom or LocusCompare and present them side by side, though it is often difficult to compare associations plotted on two different x-axes (Pruim *et al*., 2010, https://github.com/boxiangliu/locuscompare). To address this issue we built the Regional Association ComparER (RACER) package, which creates “mirror plots” for two individual associations. Mirror plots illustrate two associations, one inverted, on a shared x-axis, allowing for the direct comparison of the associated variants for two phenotypes.

## II. RACER Features

RACER was developed as a data visualization tool for the comparison of two sets of association data that share a common locus. With RACER, users can plot association data, minimally containing columns for chromosome, genomic coordinates, and p-values for an association. RACER contains a formatting function which can take any association data as input, and format it for compatibility with plotting functions. RACER also contains a function for annotating association data with population-specific linkage disequilibrium data from the 1000 genomes project using LD Link using reference SNP IDs or formatting existing linkage disequilibrium provided by the user for a specific study population (1000 Genomes Project Consortium *et al*., 2015; Machiela and Chanock, 2015). Once the association data has been formatted and annotated, RACER can produce three different types of plots: (1) a plot of a single association (**Supplemental Figure 1**), (2) a scatter plot of the p-values from two different association data sets (**Supplemental Figure 2**), or (3) a mirror plot for two associations (**Figure 1**). A vignette illustrating how to create the *MARK3* eQTL/BMD association mirror plot described below can be found at https://oliviasabik.github.io/RACERweb/articles/IntroToRACER.html.

**Figure 1.**
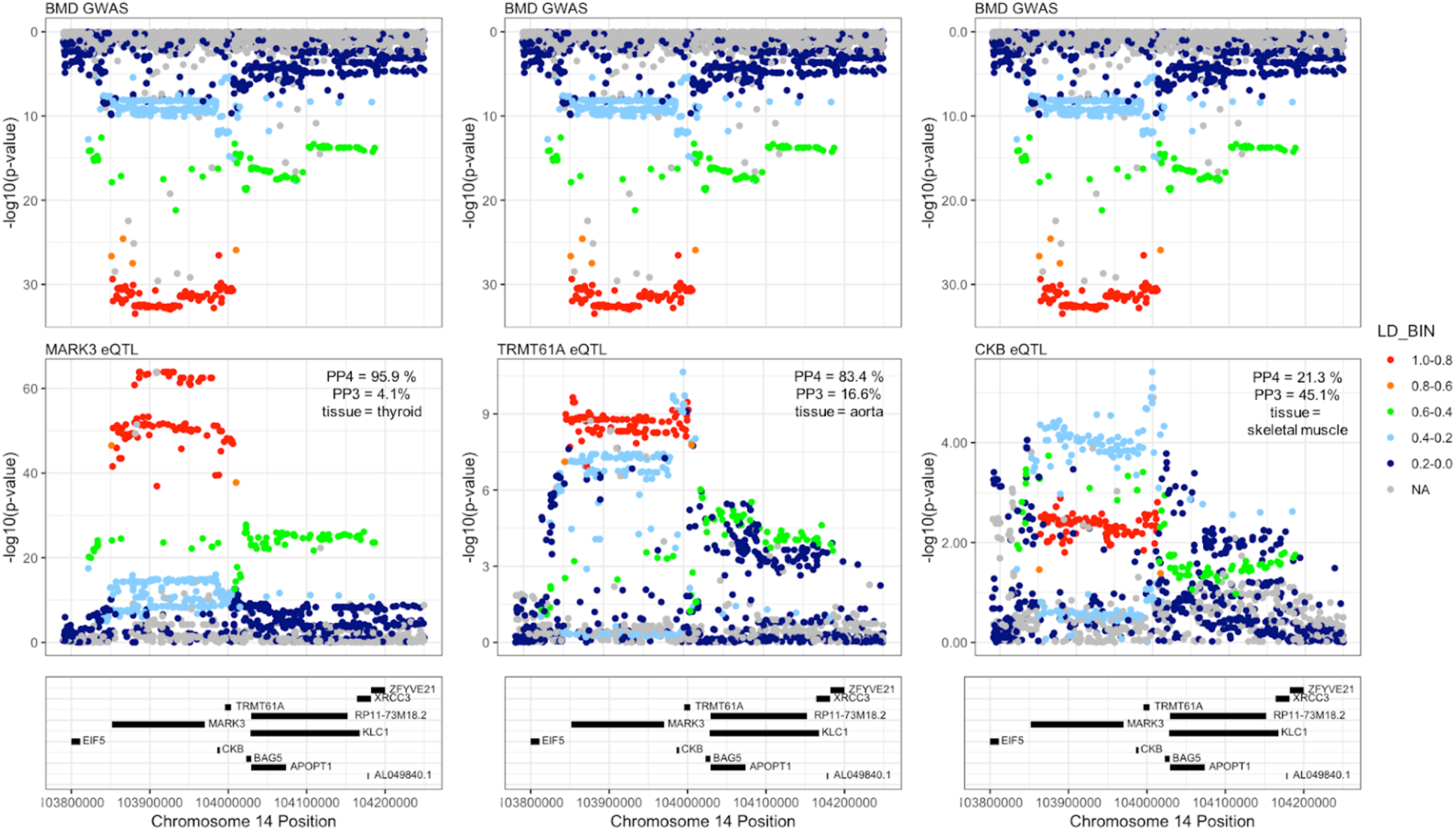
Mirror plots for *MARK3, TRMT61A* and *CKB* eQTL and a BMD GWAS locus. The mirror plots illustrate the similarity of the BMD association and *MARK3* eQTL, the complexity of the *TRMT61A* eQTL, and the dominance of the secondary association in the *CKB* eQTL.

## III. RACER Application

As a demonstration of the utility of RACER, we present a case using GTEx eQTL data to interrogate a locus on Chr. 14q32.32 associated with bone mineral density (BMD). The Chr. 14q32.32 locus spanned approximately 160 Kbp and included three genes: *MARK3, CKB* and *TRMT61A*. We previously demonstrated that the expression of all three genes were influenced by significant eQTL (p < 1.0×10^−5^) in at least one GTEx tissue (Calabrese *et al*., 2017). In the original paper, we analyzed these relationships using GTEx release v6 and BMD GWAS data from a 2012 study (Estrada *et al*., 2012). To demonstrate the use of RACER, we performed a new analysis using GTEx release v7 and BMD GWAS data from a 2017 study (**Supplemental Data 1**, **Supplemental Data 2**) (GTEx Consortium *et al*., 2017; Kemp *et al*., 2017). First, we used the coloc R package to estimate the posterior probability (PPH4) that each pair of associations were due to single causal variants (Giambartolomei *et al*., 2018). Using coloc, we observed that both *MARK3* and *TRMT61A* were likely to share a causal variant (PPH4 = 95.9% and PPH4 = 83.4%, respectively). The likelihood that the *CKB* eQTL colocalizes with the Chr. 14q32.32 locus was low (PPH4 = 21.3%)

We used RACER to create mirror plots comparing the BMD association with each of the three eQTL. This visualization of the *MARK3* and *TRMT61A* eQTL in direct comparison with the BMD association indicate that the *MARK3* eQTL has an architecture more similar to the BMD association than the *TRMT61A* eQTL. The *MARK3* eQTL is nearly identical to the BMD association; the same variants are the most significantly associated with both *MARK3* expression and BMD, and the pattern of association is similar across SNPs of decreasing LD. While the *TRMT61A* eQTL and BMD association have a PPH4>75%, which is considered sufficient evidence of a shared causal variant, it appears to be influenced by multiple associations in this region (Giambartolomei *et al*., 2018). The variants that are the most significantly associated with *TRMT61A* expression only exhibit low linkage disequilibrium with the SNPs that are the most significantly associated with BMD. However, the most significant BMD variants do see to be represented in the association, albeit at a lower level of significance. As observed in the coloc results, the *CKB* eQTL signal is dominated by an alternative signal, similar in architecture to the strongest signal in the *TRMT61A* eQTL. Using RACER, we confirmed the coloc results for *CKB* and gained a more nuanced view of the *TRMT61A* and *MARK3* results. Though this analysis does not exclude the involvement of *TRMT61A* or *CKB*, it does provide further evidence that *MARK3* is responsible for the association.

While we demonstrated the comparison between a disease association and an eQTL, RACER can be used to visualize the comparison between any two associations at a common locus, including associations for different phenotypes which may arise from a pleiotropic variant, or comparable associations arising from studies carried out in populations of different ethnicities.

## IV. Conclusion

We have developed RACER, an R package to produce mirror plots, which allow for the direct comparison of two different associations within the same locus. Mirror plots provide an effective tool for the visual exploration and presentation of the relationship between two genomic associations.

## Supporting information

## V. Acknowledgements

We would like to thank Nathan Sheffield (University of Virginia) and John Lawson (University of Virginia) for their advice in the development of RACER, and Basel Al-Barghouthi (University of Virginia), Eric Taleghani (University of Virginia), and Catherine Robertson (University of Virginia), all of whom provided critical testing and input throughout the development of RACER.

## VI. Funding

OLS was supported by a Wagner Fellowship from the University of Virginia and the University of Virginia Cell and Molecular Biology Training Grant funded by the National Institute of General Medical Sciences (T32 GM8136-31A1). This work was also supported by the National Institute of Arthritis and Musculoskeletal and Skin Diseases of the National Institutes of Health [AR071657, AR064790, and AR068345 to CRF]. The Genotype-Tissue Expression (GTEx) Project was supported by the Common Fund of the Office of the Director of the National Institutes of Health, and by NCI, NHGRI, NHLBI, NIDA, NIMH, and NINDS. Conflicts of Interest: none declared.

## Notes

#### Summary of Updates

Added middle initials to author listings

