## Supplementary material for "RACER: A data visualization strategy for exploring multiple genetic associations"

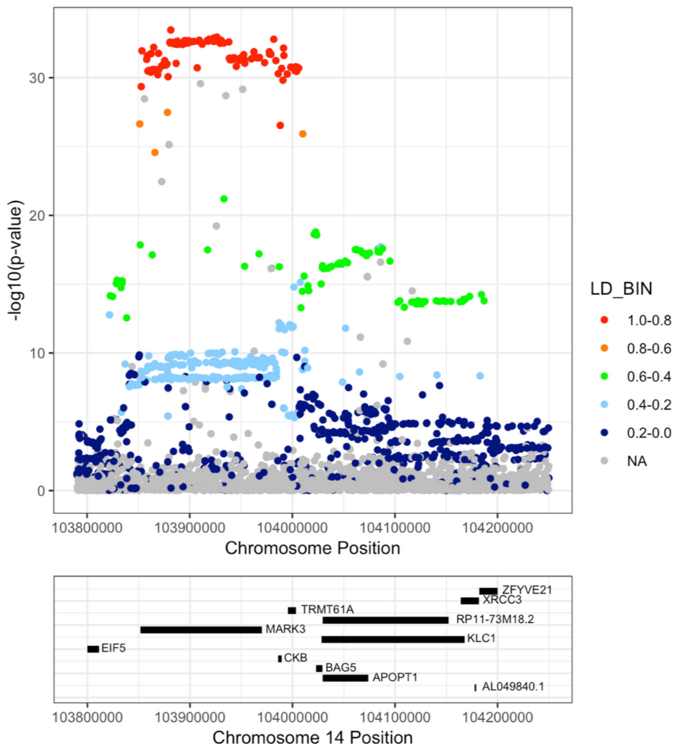


**Supplemental Figure 1. Single association plot for the BMD GWAS locus.** RACER can also be used to create a plot of a single association, for example, this plot of the Chr. 14q32.32 association for BMD.
