## Supplementary material for "RACER: A data visualization strategy for exploring multiple genetic associations"

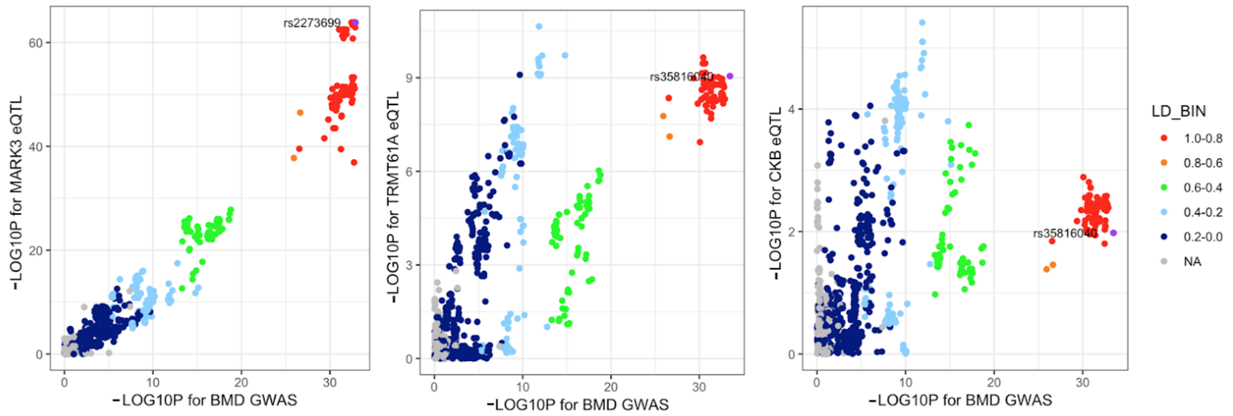


**Supplemental Figure 2. Scatter plots for *MARK3*, *TRMT61A* and *CKB* eQTL and a BMD GWAS locus.** These scatter plots illustrate the similarity of the BMD association and *MARK3* eQTL, and the complex relationship between the *CKB* eQTL, the *TRMT61A* eQTL, and the BMD association.
