## Supplementary material for "RACER: A data visualization strategy for exploring multiple genetic associations"

### Supplemental Data Manifest

I. Supplemental Data
   A. Supplemental Data 1. Example BMD GWAS association data from the Chr. 14q32.32 locus.
   B. Supplemental Data 2. Example GTEx eQTL data for *MARK3* from thyroid.

II. Supplemental Figures
   A. Supplemental Figure 1. Single association plot for the BMD GWAS locus.
   B. Supplemental Figure 2. Scatter plots for *MARK3*, *TRMT61A* and *CKB* eQTL and a BMD GWAS locus.
